# Eukaryotic rather than prokaryotic microbiomes change over seasons in rewetted fen peatlands

**DOI:** 10.1101/2020.02.16.951285

**Authors:** Haitao Wang, Micha Weil, Kenneth Dumack, Dominik Zak, Diana Münch, Anke Günther, Gerald Jurasinski, Gesche Blume-Werry, Jürgen Kreyling, Tim Urich

## Abstract

In the last decades, rewetting of drained peatlands is on the rise worldwide, to restore the significant carbon sink function. Rewetted peatlands differ substantially from their pristine counterparts and can, thus, be considered as novel ecosystems. Despite the increasing understanding of peat microbiomes, little is known about the seasonal dynamics and network interactions of the microbial communities in these novel ecosystems, especially in rewetted groundwater-fed peatlands, i.e. fens. Here, we investigated the seasonal dynamics in both prokaryotic and eukaryotic microbiomes in three common types of fens in Northern Germany, namely percolation fen, alder forest and coastal fen. The eukaryotic microbiomes, including fungi, protists and metazoa, showed significant changes of their community structures across the seasons in contrast to largely unaffected prokaryotic microbiomes. The co-occurrence network in the summer showed a distinct topology compared to networks in the other seasons, which was driven by the increased connections among protists, as well as between protists and the other microbial groups. Our results also indicated that the dynamics in eukaryotic microbiomes differed between fen types, specifically in terms of saprotrophs, arbuscular mycorrhiza and grazers of bacteria. Our study provides the insight that microbial eukaryotes mainly define the seasonal dynamics of microbiomes in rewetted fen peatlands. Accordingly, future research should unravel the importance of eukaryotes for biogeochemical processes, especially the under-characterized protists and metazoa, in these novel yet poorly understood ecosystems.

## 1. Introduction

Peatlands store over 30% of the earth’s soil carbon although they only cover nearly 3% of the global land area (Gorham 1991; Hugelius et al., 2020). Water logging contributes to the high level of soil organic carbon since peat, consisting of incompletely decomposed organic matter derived mostly from plant residues, accumulates when the oxygen is deficient (Page and Baird 2016). However, in some countries more than 90% of peatlands were drained for agriculture including grasslands, resulting in significant carbon loss through the emission of greenhouse gases and thus contributing to global warming (Haddaway et al., 2014; Kasimir‐Klemedtsson et al., 1997; Lamers et al., 2015). Fens are base-rich and groundwater-fed peatlands and the dominant peatland type in temperate zones. The conversion of natural fens to agricultural lands also leads to biodiversity loss due to intensive dehydration, tillage and fertilization (Lamers et al., 2015). Rewetting of drained peatlands re-establishes water-logged and thus anoxic conditions, then the existing peat carbon pool is protected from decomposition through a variety of processes and new carbon may be stored. However, it remains unclear to what extent the ecological functioning of drained peatlands recovers following rewetting. Since previous agricultural management of these peatlands may severely and permanently alter the environmental conditions, rewetted peatlands are not pristine anymore and thus are suggested to be novel ecosystems (Jurasinski et al., 2020b).

The microbiome, a major driver of biogeochemical cycling, plays a pivotal role in peatland ecosystems. The composition of prokaryotic microbiomes in general were significantly shaped by rewetting (Emsens et al., 2020; Weil et al., 2020), associated with an increased relative abundance of anerobic microbial groups in rewetted fens (Weil et al., 2020). Studies on rewetted peatlands also focus on the methane emissions and associated methanogens (Urbanová and Bárta 2020; Wen et al., 2018), as increased methane emissions are a major concern after rewetting. Despite of the rise in the research interest on peat microbiomes, the majority of the ecological roles of microbiomes still remains under-characterized in rewetted peatlands, especially in fen peatlands whose pristine states are not even well understood (Jurasinski et al., 2020b).

One such under-estimated aspect is the eukaryotic fraction of microbiomes which is also in general underappreciated. Fungi are major cause of plant decomposition, which eventually leads to the release of CO_2_ to the atmosphere (Bani et al., 2018). This so called soil respiration is a crucial element of global carbon cycling (Van Geffen et al., 2010). Protists constitute the majority of microbial eukaryotes and play key roles in microbial foodwebs, especially as predators of bacteria and fungi (Geisen et al., 2018). Bacterivorous protists significantly shape the composition of the soil bacterial community and make nutrients available for plants (Geisen et al., 2018; Ronn et al., 2002). Some protists are also important primary producers, while others represent important parasites (Geisen et al., 2018). Accordingly, the functions of protists are manifold. Microbial metazoa are the most ignored group of eukaryotes in microbial ecology studies (Bik 2019). Nematodes, the most abundant animals worldwide, fill all trophic levels in soil foodwebs and thus play critical roles in regulating carbon and nutrient dynamics in soils (Bardgett and van der Putten 2014; van den Hoogen et al., 2019). However, to date, there is only little information available on eukaryotic groups in rewetted fens, particularly protists and metazoa.

Unraveling the dynamics of community composition together with relevant environmental factors may help to carve out predictable spatial and temporal patterns (Fuhrman et al., 2015) and, thus, further our understanding of the ecological consequences resulting from these patterns. A limited number of studies have investigated the spatial variability of microbiomes in rewetted fens (Emsens et al., 2020; Weil et al., 2020), while there is no study yet on seasonal dynamics. Moreover, it has been shown that environmental characteristics including soil salinity, nitrogen and phosphorus concentrations, water table, as well as water and nutrient fluxes follow seasonal dynamics in peatlands (Feng et al., 2020; Kennedy et al., 2018; Kieckbusch and Schrautzer 2007; Walter et al., 2018). These environmental factors certainly interplay with yet unexplored seasonal dynamics of peat microbiomes (Liu et al., 2012; Santoro et al., 2006; Wang et al., 2018; Weil et al., 2020). Thus, understanding the seasonal dynamics in the microbiomes in rewetted fens could improve the understanding of ecosystem functionality and resilience after rewetting.

About two decades ago, a peatland restoration program was initiated in the state of Mecklenburg-Vorpommern (M-V) in Northern Germany, and in this course more than 20,000 ha of drained peat-soils were rewetted (Zerbe et al., 2013). In this region, the relatively high number of rewetted sites in different peatland types provides an excellent opportunity to study the influence of rewetting on ecosystem dynamics and functioning. In a previous study, we observed that rewetting significantly changed the community compositions of both prokaryotes and eukaryotes in the three most relevant peatland types of this region, namely alder forest, coastal fen and percolation fen (Weil et al., 2020), but it remained unclear if and how these dynamics change over seasons. Here, in this follow-up study, we focus on the seasonal dynamics in both prokaryotic and eukaryotic microbiomes in the three types of fens by including three more time points covering a whole year. We mainly focused our analyses on the rewetted sites, but comparisons between drained and rewetted sites were made when necessary to add to our understanding of the involved processes. We characterized the seasonal dynamics in prokaryotic and eukaryotic microbiomes as well as their network interactions, with a special focus on fungi, protists and metazoa. We aim to understand the temporal changes of microbiomes per site as well as with varied peatland conditions across sites, which is among the high priority questions in understanding the resilience of peatland ecosystem (Ritson et al., 2021). Eventually, we aim to provide the first insight into the seasonal dynamics of microbiomes, including microbial eukaryotes, in these novel yet poorly understood ecosystems.

## 2. Material and methods

### 2.1. Study sites and soil sampling

The six study sites are located in Mecklenburg-Vorpommern (MV) in Northeastern Germany (Fig. S1) and cover the three major peatland types (alder forest, coastal fen and percolation fen) of the region. For each peatland type, two sites were selected. All sites were subject to a decade-long drainage in the past but each one of the two sites per peatland type was subsequently rewetted over a decade ago. The rewetted sites are named as PW (rewetted percolation fen), AW (rewetted alder forest) and CW (rewetted coastal fen), while their drained counterparts are named accordingly as PD (drained percolation fen), AD (drained alder forest) and CD (drained coastal fen) (Fig. S1). The six study sites were selected to be representative of the respective fen types regarding basic site characteristics (Jurasinski *et al.*, 2020; Schwieger *et al.*, 2020). Spatial replicates per site can be seen as independent and representative for the respective vegetation type as spatial auto-correlation is almost absent in these peatlands (Koch & Jurasinski, 2015; Koch *et al.*, 2014). The details regarding the history, management and characteristics of these six sites are described in (Jurasinski et al., 2020a).

The peat soils were sampled from these six sites in April 2017 (spring), August 2017 (summer), November 2017 (autumn) and February 2018 (winter). At each site, samples were taken at three spots as replicates at depths of 5-10 cm, 15-20 cm and 25-30 cm from the top soils. Samples were taken using a soil core and then stored in sterile Nasco Whirl-PAK baggies. The collected soils were immediately blended and transported to the laboratory with an ice box. Soils were processed on the same or the next day of sampling. In total, 216 samples (3 depths × 6 sites × 4 seasons × 3 replicates) were collected.

### 2.2. Soil edaphic properties

Soils moisture was gravimetrically measured by drying the soil over night at 90 °C until mass constancy. The concentrations of dissolved organic matter (DOM), soluble reactive phosphorus (P), N-NH_4_^+^ and N-NO_x_^−^ were measured as previously described (Weil et al., 2020). The detected DOM was categorized into three groups: (i) biopolymers (BP-S), (ii) humic or humic-like substances including building blocks (HL-S), and (iii) low molecular-weight substances (LM-S), according to (Heinz and Zak 2018). The pH and salinity of water samples taken from wells installed at each study site were measured manually by digital meters in April 2017. The data of the other three seasons were manually measured at groundwater wells with Aquaread water sensors (AP-2000/AP-2000-D). Water levels and soil temperatures were also continuously monitored at each site using Campbell Scientific CR300 Dataloggers (Logan, USA) and HOBO Dataloggers (Bourne, USA), respectively.

### 2.3. DNA extraction and sequencing

DNA was extracted from 0.25 g soil using the DNeasy PowerSoil Kit (QIAGEN, Hilden, Germany) according to the manufacturer’s instructions with some modification. Vortex in the bead beating step was replaced with a FastPrep^®^-24 5G instrument (MP Biomedicals, Santa Ana, USA), with an intensity of 5.0 m/s for 45 s. The extracted DNA samples were sent to LGC Genomics GmbH (Berlin, Germany) for 16S rRNA and 18S rRNA gene amplicon sequencing, with Illumina Miseq 300 bp paired-end platform. Information on primers used is described in the previous study (Weil et al., 2020). All the sequencing data were deposited to the European Nucleotide Archive of EMBL (European Molecular Biology Laboratory). The study accession number is PRJEB36764.

### 2.4. Sequencing data analysis

16S rRNA and 18S rRNA gene amplicon sequences were processed separately. For each processing, the raw sequence reads were demultiplexed with barcodes, adapters and primers removed using the Illumina *bcl2fastq* software. The data were then processed with the *dada2* (v1.8.0) pipeline (Callahan et al., 2016) in R v3.6.3. The qualities of the sequences were checked, and sequences failing to meet the filter scores (maxEE=2, truncQ=2, maxN=0) were discarded. The filtered sequences were de-replicated and clustered into ASVs, and paired-end sequences were merged. The chimeric sequences were then de-novo checked and removed. The final representative sequence of each ASV was assigned to taxonomy against a modified version of the SILVA SSUref_NR_128 database (Lanzén et al., 2012). ASVs with only one sequence were removed. ASVs of 16S rRNA amplicons that were assigned to chloroplast or mitochondria were also removed. Several samples were discarded due to their low sequence numbers (<1,000), resulting in a total of 209 samples for further analyses.

To mitigate the impact of uneven sequencing depths, ASV count per sample was rarefied to the minimum sequence number among samples for alpha diversity (Shannon index) estimation. For beta diversity, the tables of ASV counts were normalized using metagenomeSeq’s CSS (Paulson et al., 2013), and the community compositions were analyzed using non-metric multidimensional scaling (NMDS) based on Bray-Curtis dissimilarity distances. The 16S rRNA taxonomic profiles were converted to putative functional profiles using FAPROTAX which maps prokaryotic clades to established metabolic or other ecologically relevant functions (Louca et al., 2016). However, some modifications were made. The ASVs involved in nitrification were further verified by blasting against the NCBI database following (Arce et al., 2018). The anaerobic methanotrophic groups ANME were excluded from methanogens. Similarly, the fungal taxonomic profiles were also converted to ecological guilds using FUNGuild (Nguyen et al., 2016). The assigned functional traits of fungi mainly comprise of different saprotrophic types, arbuscular mycorrhiza, ectomycorrhiza, endophytes and pathogens of plants and animals. Functional traits of protists and metazoa were assigned according to (Adl et al., 2019; Dumack et al., 2020; Yeates et al., 1993). The assigned functional traits of protists comprise of plant and animal parasites, saprotrophs, eukaryvores, bacterivores, omnivores, algae and mixotrophs. The assigned functional traits of metazoa consist of fungivores, omnivores, predators, animal parasites, substrate feeders, filter feeders, and plant and bacteria feeding organisms.

Co-occurrence networks were constructed to explore the seasonal dynamics of potential interactions between species, including prokaryotes, fungi, protists and metazoa. As there were more rare species and uneven sequencing depths in prokaryotic data, they were rarefied to 1,000 sequences per sample. Then all ASV counts were normalized with the aforementioned method. Each network regarding one season was based on 51 to 53 communities. Only ASVs that occurred in at least eight communities (15% of the samples) were kept for network analysis. The pairwise Spearman’s rank correlations were performed with *Hmisc* package (Harrell Jr and Harrell Jr 2019) and all *P*-values were adjusted by the Benjamini and Hochberg false discovery rate (FDR) method. The cutoffs of correlation coefficients and adjusted *P*-values were 0.7 and 0.01, respectively. The significant correlations were visualized using *igraph* package (Csardi and Nepusz 2006). Networks consist of modules that are intensively connected to themselves but sparsely connected to other modules. The modularity was identified from each network via greedy optimization using *igraph*. Then the within- and among-module connectivities were calculated as Zi (within-module degree) and Pi (participation coefficient) values, respectively, according to (Guimera and Nunes Amaral 2005). The nodes in the networks were further categorized into module hubs (Zi > 2.5; Pi < 0.62), connectors (Zi < 2.5; Pi > 0.62), network hubs (Zi > 2.5; Pi > 0.62) and peripherals (Zi < 2.5; Pi < 0.62) referring to their topological roles (Poudel et al., 2016).

### 2.5. Statistical analysis

The statistical analyses were done with R 3.6.3. The significance of the seasonal dynamics of soil edaphic parameters and predicted microbial functions was identified with the non-parametric Kruskal-Wallis test provided in the *vegan* package (Oksanen et al., 2013). The ASVs that showed significant seasonal dynamics were also identified with Kruskal-Wallis test. Permutational Multivariate Analysis of Variance (PERMANOVA) was performed to test the significance of seasonal dynamics in prokaryotic and eukaryotic community compositions using the *vegan* package. Venn diagrams showing the number of shared differentially abundant ASVs between sites were plotted using the *VennDiagram* package (Chen and Boutros 2011). All *P*-values for multiple comparisons were adjusted by the FDR method and the null hypothesis was rejected for (adjusted) *P* < 0.05.

## 3. Results and discussion

### 3.1. Microbiome structures of prokaryotes and eukaryotes in all sites

The 16S rRNA and 18S rRNA sequences were clustered into 25,864 ASVs and 15,937 ASVs, respectively. The eukaryotic ASVs were further categorized into four groups: plants, fungi, protists and metazoa. The protists are defined as all eukaryotes excluding plants, fungi and animals. To verify if the pattern of community structures regarding water status (drained and rewetted) and peatland type (percolation fen, alder forest and coastal fen) we observed before (Weil et al., 2020) stayed stable over the seasons, we constructed NMDS plots based on all six sites (Fig. S2). Even with now more time points, the NMDS plots still showed the prokaryotic and eukaryotic community compositions to be significantly driven by both, water status and peatland type. This is in line with (Weil et al., 2020), suggesting that rewetting affects microbial communities by magnitudes larger than seasonality. Since the communities in these six sites were distinctly different, in the following we focus on the seasonal dynamics in each site.

### 3.2. Seasonal dynamics in soil conditions

We measured edaphic properties to evaluate the seasonality of the environmental conditions in the studied fens. The soils underwent significant seasonal dynamics in all three rewetted sites (Fig. 1). The DOM changed significantly over the seasons in all rewetted sites, with a remarkable increase in autumn resulting from changes of the three components (BP-S, HL-S and LM-S) all showing the same trend. The water table was stable and high in the three rewetted sites until April 2018 (Fig. S3), and the water content also showed no significant changes over seasons (Fig. 1). In AW, the NH_4_^+^ concentration was significantly lower in spring than in the other seasons, while the NO_x_^−^concentration showed an opposite pattern. P concentration and salinity showed significant dynamics in AW and CW. Salinity was much higher and showed more variations in CW due to seawater flooding. In summer, there was an increase of salinity in CW, presumably driven by a combination of increased evaporation and water uptake by plants. Since pH was measured with different methods on spring samples and during the other sampling campaigns, we excluded spring from our comparisons. The three seasons witnessed significant changes of pH in all rewetted sites. pH was generally lower in summer compared with autumn and winter. The soil temperature followed a clear seasonal pattern from August 2017 to July 2018 (Fig. S4). Moreover, the drained sites also showed many significant dynamics of these properties, but with some different patterns (Fig. S5). Thus, the soil conditions varied significantly across the seasons, which could drive the seasonal dynamics we observed in the microbiomes.

**Fig. 1.**
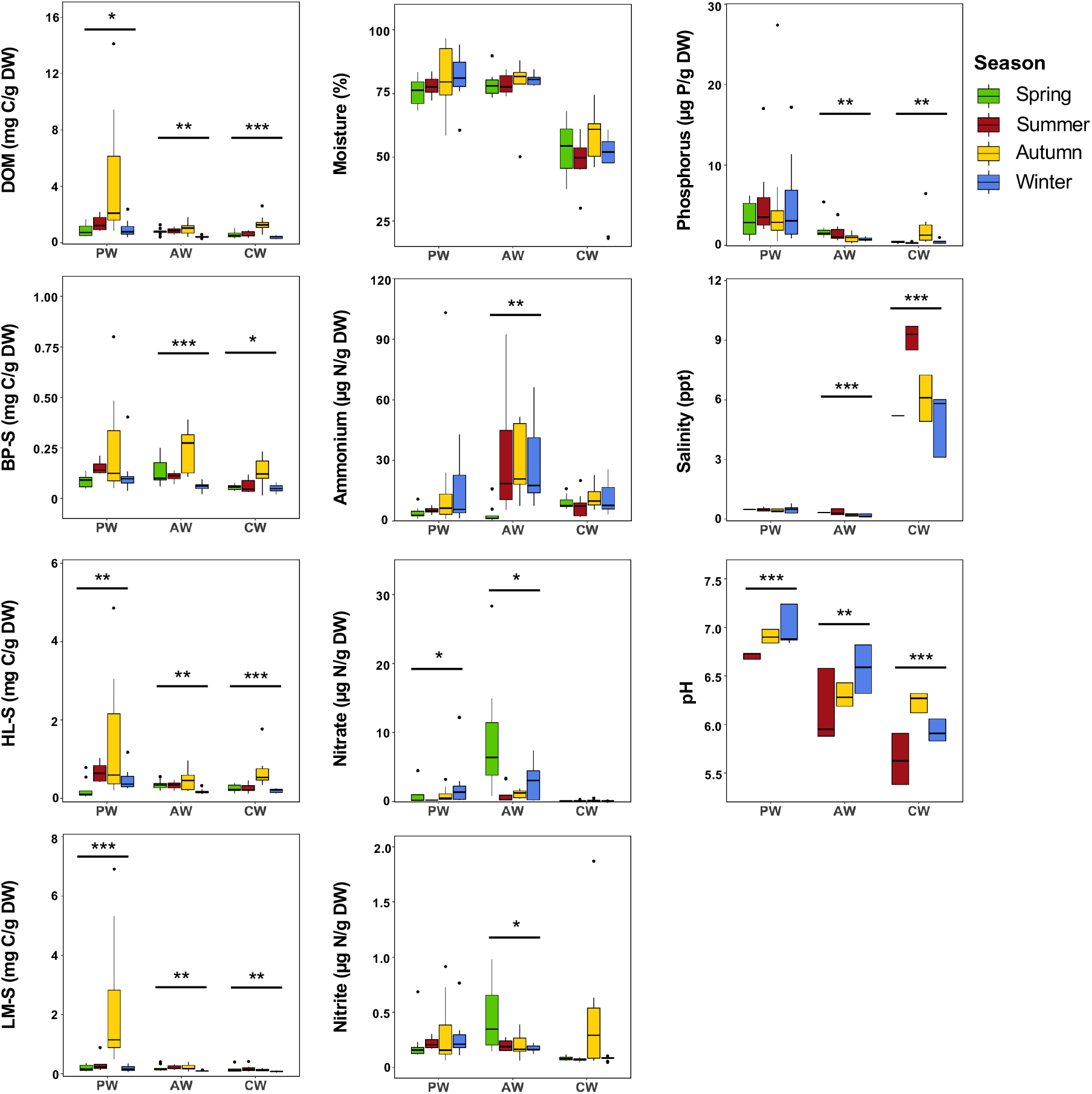
Seasonal dynamics in soil edaphic properties in the three rewetted sites. Asterisks indicate the significant seasonal changes in one site (Kruskal-Wallis test, *adjusted *P*<0.05, **adjusted *P*<0.01, ***adjusted *P*<0.001). DOM, dissolved organic matter; BP-S, biopolymer substances; HL-S, humic-like substances; LM-S, low-molecular substances; DW, dry weight. pH in spring is not shown as a different measure method was used in spring.

### 3.3. Seasonal dynamics in prokaryotic microbiomes

The alpha diversity of the prokaryotic microbiome showed significant seasonal dynamics in PD and CD, while no significant dynamics were observed in any of the rewetted sites (Fig. S6 and S7). NMDS plots based on ASV levels in each site showed the four season-clusters (Fig. 2 and S8). Only in AW the seasonal dynamics of the prokaryotic communities showed significant changes, according to PERMANOVA (Table S1). At the phylum or class level, the variations of a few taxa across the seasons were observed in AW, while the taxa in the other sites stayed rather stable (Fig. S9 and S10) suggesting only minor seasonal dynamics in prokaryotic microbiomes in the studied fen peatlands. We also investigated the seasonal dynamics at a functional level using the functional guilds predicted by FAPROTAX. We focused on eight dominant and representative functional guilds regarding carbon and nitrogen cycling, and decomposition (Fig. S11 and S12). Our results show that the seasonal dynamics of these functional guilds are weak in both, rewetted (Fig. S11) and drained sites (Fig. S12), which also indicates a lack of seasonal dynamics in prokaryotes.

**Fig. 2.**
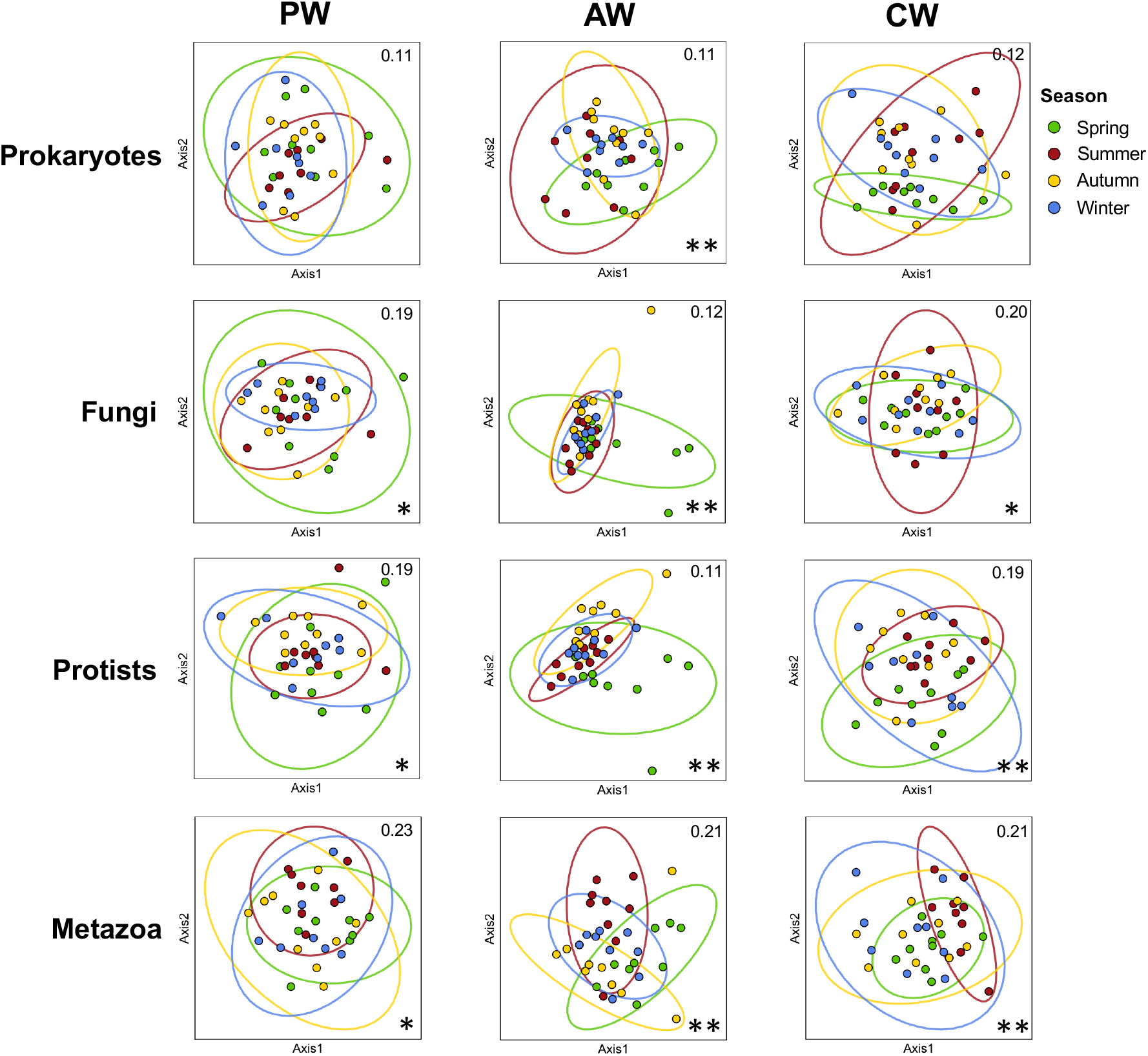
NMDS plots based on the Bray-Cutis dissimilarities showing community compositions of prokaryotes and eukaryotes (fungi, protists and metazoa) in each rewetted site. Numbers on the top right indicate the stress values. Asterisks represent the significance level of PERMANOVA (*adjusted *P*<0.05 or **adjusted *P*<0.01) as indicated in Table S1.

Contrastingly, most of the assessed soil parameters, including soil temperature, the most important indicator of seasonality, showed significant seasonal changes (Fig. 1 and S3) suggesting that peat prokaryotic communities do not react strongly to weather-driven temporal changes of environmental conditions. This resilience to external forces such as temperature and nutrient supply might result from the internal feedback mechanisms, including competition, viral infection, and predator-prey interlinkages, maintaining a relatively stable state of community in seasonally changing environments (Fuhrman et al., 2015). However, the water content (Fig. 1) and the water level within the studying period (from September 2017 to Feburary 2018, Fig. S3) showed no obvious seasonal patterns, although this could have been expected in such a system. The stable water status among seasons might have stablized the prokaryotic communities, since water-saturation or high-water levels may create homogeneity and weak niche differentiation that leads to stronger interactions between microbes (Faust and Raes 2012), resulting in high resilience to environmental stresses. However, one year observation might be a limited basis for understanding seasonal patterns. In addition, we experienced quite unusual weather patterns over the course of our study period with very wet end of 2017 followed by an exceptionally dry summer in 2018 (Fig. S3). Therefore, a longer-term monitoring of prokaryotic dynamics in peatlands is needed in future studies.

### 3.4. Seasonal dynamics in eukaryotic microbiomes

In stark contrast to the prokaryotic microbiomes, the eukaryotic microbiomes of fungi, protists and metazoa all showed significant seasonal dynamics in all sites (Fig. 2 and Table S1), even if their overall diversity did not change strongly across seasons (Fig. S6 and S7). This indicates that eukaryotic communities are more sensitive to environmental changes compared to prokaryotic communities. Supporting this, Zhao et al., (2019) found that protist communities were more affected by the application of nitrogen fertilizer than bacterial and fungal communities in diverse agricultural soils. Zhao *et al.*, (2019) also suggest that protist communities, showing the strongest seasonal dynamics, serve as the most sensitive bio-indicators of soil changes. Oshima et al., (2020) reported that protists and fungal communities were less resistant and resilient to high temperatures than the bacterial community in soils. The kinetic stability, a crucial property of proteins for the adaptation and survival of microorganisms, is prevalent in prokaryotes but uncommon in eukaryotes (Xia et al., 2010), which might explain the higher sensitivity of eukaryotes to environmental changes. As our findings suggest that eukaryotes mainly define the seasonal dynamics in microbiomes in rewetted fen peatlands, in the following seasonal dynamics in the three major eukaryotic groups are discussed in detail as well as major microbial drivers of these dynamics.

#### 3.4.1. Seasonal dynamics in fungal communities

Ascomycota predominated the fungal community in PW and all fungal phyla in this site showed only slight changes over the seasons (Fig. S9). In AW, both Ascomycota and Basidiomycota were dominant. Ascomycota showed lower relative abundance in summer and autumn, while Basidiomycota changed abundances in opposite direction. Interestingly, the arbuscular mycorrhiza Glomeromycota constituted a major part of the fungal community in CW. They were more abundant in summer and autumn, which is opposite to the changes in Ascomycota observed in CW.

To identify the key species who were driving these seasonal dynamics, we used Kruskal-Wallis test to determine the ASVs that were significantly different among the seasons. The full list of these ASVs and their associated taxonomies and assigned functional traits is provided in Table S3. The differentially abundant ASVs were different in the three sites, since they were barely shared between sites for all three groups (Fig. 3). The single most dominant fungal ASV in the freshwater sites (PW and AW) was ASV11, a saprotrophic fungus that showed the highest abundance in spring and then decreased consistently until winter (Fig. 3). As degradation of the plant residues is the major process of decomposition in peatlands (Page and Baird 2016), this fungus probably represents a plant saprotroph involved in plant decomposition. The increased decomposition in spring and summer might lead to an increased accumulation of available organic carbon, which may contribute to the significant increase of DOM in autumn (Fig. 1).

**Fig. 3.**
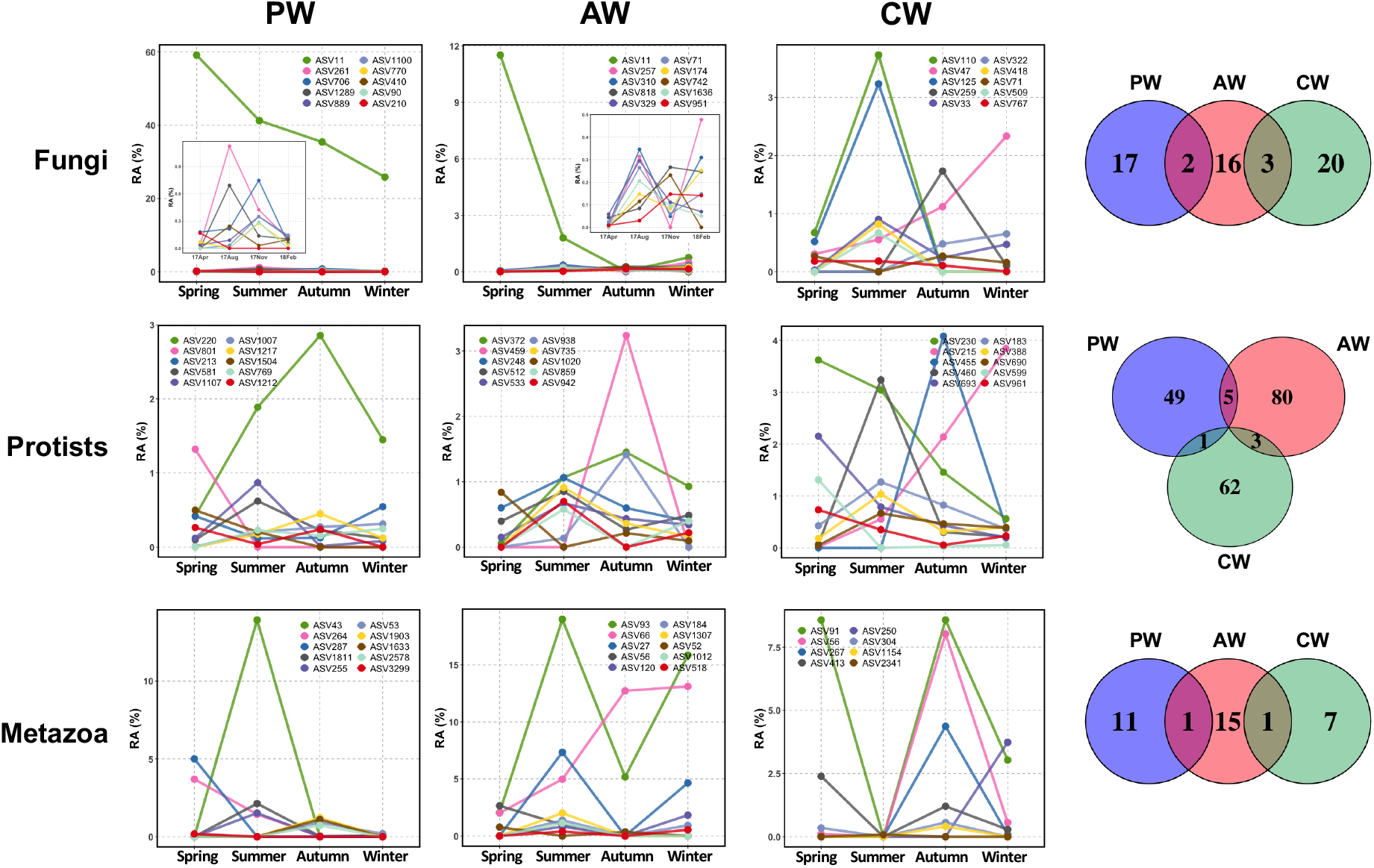
Changes of relative abundances of eukaryotic ASVs showing significant seasonal dynamics. Only the most abundant ASVs in each site are shown in the line charts. The Venn diagrams show the numbers of shared and unique ASVs (showing significant seasonal dynamics) among the three rewetted sites.

The coastal site (CW), however, showed strongly different patterns. The major microbial drivers here were two abundant arbuscular mycorrhiza (ASV110 and ASV125), which sharply increased in summer, in line with the pattern observed for the whole Glomeromycota (Fig. S9). Plants control mycorrhizal colonisaton and activity, so this pattern is to be expected with plant activity patterns during the growing season (Gavito and Varela 1993; Johnson et al., 1992). Additionally, the increase in temperature positively contributes to the growth and functions of arbuscular mycorrhiza fungi (Heinemeyer and Fitter 2004; Kytoviita and Ruotsalainen 2007). Land and Schönbeck (1991) found that, in Northern Germany, the mycorrhizal colonization in three different soil types increased from April until July with a temperature range between 0 °C and 20 °C, which is similar to the range of temperatures in our study (Fig. S4). The congruent results between this study and our study suggest the general response of arbuscular mycorrhiza to temperature changes in this region.

The divergent compositional changes of fungal communities between freshwater and coastal sites are likely caused by the different salinity conditions. The intrusion of sea water in the coastal site resulted in a higher salinity (Fig. 1) which may inhibit the growth of saprotrophic fungi and thus the decomposition, as observed in soils and sediments in general (Qu et al., 2018; Wichern et al., 2006). Furthermore, there is evidence that arbuscular mycorrhiza alleviate the salinity stress on plant growth and nutrient uptake (Daei et al., 2009; Porcel et al., 2011), hinting at a tolerance of mycorrhizal fungi to saline conditions. In such habitats, the plants depend more strongly on mycorrhiza as they need them to avoid salt stress (Dastogeer et al., 2020), which might be the reason for the high relative abundance of arbuscular mycorrhiza in the coastal site (Fig. S9).

#### 3.4.2. Seasonal dynamics in protist communities

The protist communities were dominated by Apicomplexa, Cercozoa, Chlorophyta, and Ciliphora in the freshwater sites PW and AW, while Labyrinthulomycetes were additionally abundant exclusively in CW (Fig. S9). The major changes in PW included a decrease of Ciliphora in summer and an increase of Cercozoa in autumn. In CW, Cercozoa decreased in summer while Kinetoplastea and Labyrinthulomycetes increased in summer.

Bacterivores accounted for the majority of seasonally changing microorganisms in our study (Fig. 3 and Table S3). These phagotrophic protists are the principal consumers of bacteria in almost all environments leading to the release of nutrients (Fenchel 2013; Geisen et al., 2018). Their higher relative abundance in summer suggested that they might play an important role in trophic interactions in summer. However, the little seasonal variation in prokaryotes indicated that the variations in protists might not be from trophic interactions but likely be driven by changes in abiotic conditions. The warmer temperature (15-20 °C) in summer than in the other seasons (Fig. S4) might contribute to this, since the growth rate of bacterivorous protists showed positive relationships with temperature within the range of 0-40 °C (Rose and Caron 2007). Interestingly, many plant parasites showed their highest abundance in autumn at the end of the growing season when the TOM was also the highest (Fig. 1, 3 and Table S3), suggesting the potential impact of plant parasites on the release of organic matter, or *vice versa*. This might also be related to natural plant senescence at this time of the year which needs further investigation.

#### 3.4.3. Seasonal dynamics in metazoan communities

Arthropoda and Nematoda were the dominant metazoan phyla in all three sites (Fig. S9). In PW, there was a sharp increase of Platyhelminthes in summer and increases of Arthropoda and Gastrotricha in autumn, while Annelida decreased in summer and autumn. In AW, there were sharp increases of Annelida and Arthropoda and a sharp decrease of Nematoda in summer. In CW, Arthropoda consistently increased from spring onwards, but Nematoda showed an opposite change. Annelida and Rotifera only occurred in autumn.

The seasonally changing metazoa showed different seasonal patterns in the three sites (Fig. 3), suggesting that the different environmental conditions of the sites impacted their dynamics. The omnivorous and bacterivorous metazoa were the major drivers of seasonal dynamics in PW, while only bacterivorous metazoa accounted for the majority in AW (Table S3). Interestingly, almost all metazoan drivers with significant temporal dynamics in CW were omnivores (Table S3). However, the existence of some highly abundant protist omnivores among the drivers in AW and many protist bacterivores among the drivers in CW compensated for the lack of metazoan omnivorous drivers in AW and the lack of metazoan bacteria feeding drivers in CW, respectively. This suggests a kind of balance between protists and metazoa on grazing of bacteria and small animals, which might stabilize the soil foodwebs in different environments.

### 3.5. Seasonal dynamics in network interactions

Correlations in a network may indicate potential interactions among different microbial groups (Faust and Raes 2012). The relationships between prokaryotes and different groups of eukaryotes in ecological networks are fundamental to maintain stable foodwebs. Eukaryotic microbiomes showed strong seasonal dynamics and whether their dynamics had impact on the network interactions is of great interest. We therefore constructed networks based on the four seasons to show the links between prokaryotes, fungi, protists, and metazoa (Fig. 4).

**Fig. 4.**
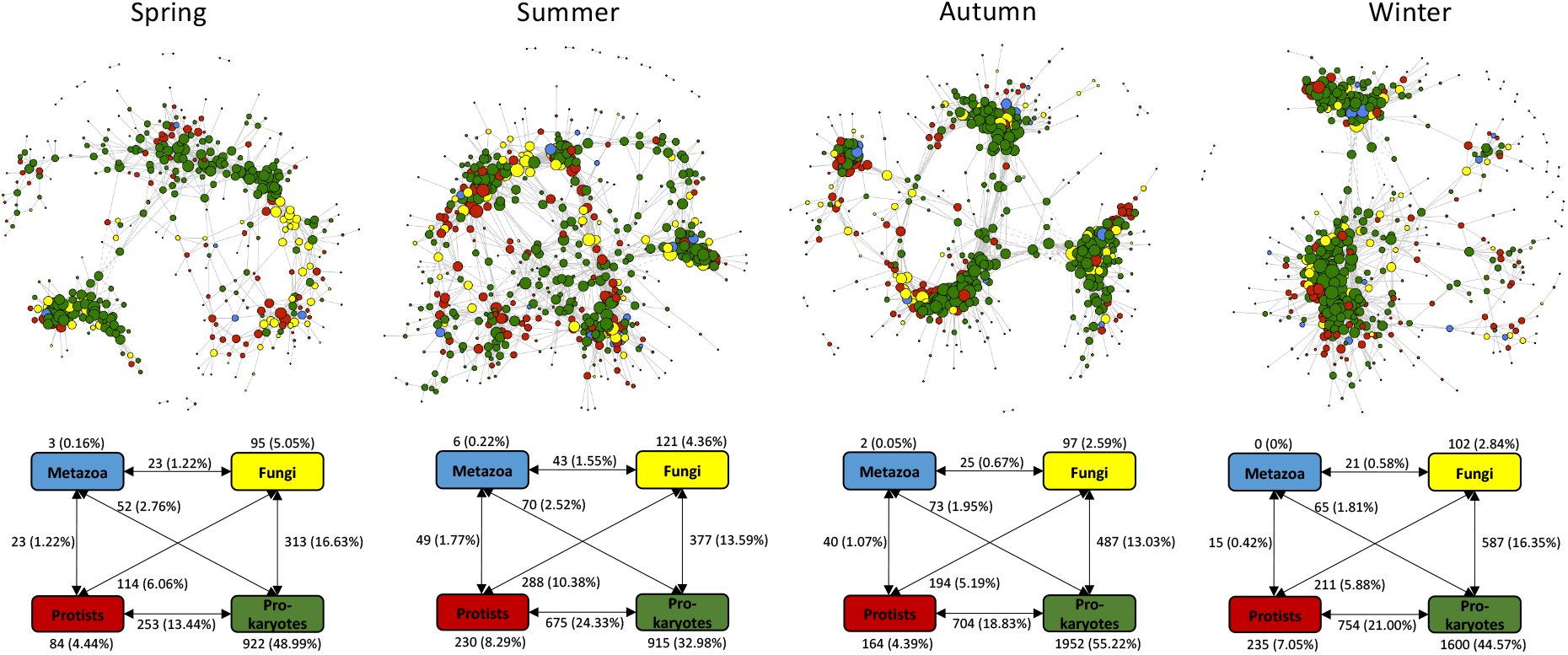
Co-occurrence networks showing seasonal dynamics in potential interactions between prokaryotes and eukaryotes (fungi, protists and metazoa). Number between two boxes indicates the number of connections between these two groups of microbes. Ratio of the edges to total edges in this network is shown in the brackets. Solid lines in the network indicate positive correlations while dashed lines indicate negative correlations. The sizes of the nodes are scaled to the degrees of the nodes.

The four constructed networks show different topological features (Table S2). The network of spring had a lower complexity (less edges and vertices) compared to the other networks. However, the higher average path, assortativity degree and diameter in spring indicated a more dispersed and even topology of the network in spring. Among all the networks, the edges among prokaryotes accounted for the largest proportion of the total edges, followed by edges between prokaryotes and protists or fungi, while metazoa only contributed to a small proportion of the edges (Fig. 4). Two distinct major clusters were observed in spring, autumn and winter, while they disappeared in summer (Fig. 4), indicating different interactions of the taxa in summer. This was mainly due to the increased edges among protists, and between protists and other taxa. The edges among protists accounted for 8.29% in summer, which was nearly doubled compared to spring and autumn and was slightly higher than winter (Fig. 4). The edges between protists and prokaryotes also increased significantly compared to spring and to a lesser extent compared to autumn and winter, accounting for 24.33% (Fig. 4). The edges between protists and fungi, which accounted for 10.38% of all edges, were also nearly doubled compared to the other seasons, while the edges between protists and metazoa showed less increase (Fig. 4). Remarkably, the edges among prokaryotes in summer were significantly reduced, which might be due to the grazing pressure by the protists in summer when we also observed some significant increases of relative abundances of protist bacterivores (Fig. 3). These results indicate that protist communities have a stronger effect on driving the seasonal dynamics in the network interactions compared to the other microbial communities.

We obtained many module hubs from each season, while only several connectors were found in summer and winter (Fig. S13). The full list of the module hubs and connectors regarding their taxonomy and functional traits were provided in Table S4. In spring, many prokaryotes including two AOB and one wood-saprotrophic fungus were identified as module hubs. However, the majority of the hubs in summer were eukaryotes, including several saprotrophic fungi, one pathogenic fungus, several algal and plant-parasitic protists and one plant feeding metazoa. In autumn, the modules hubs consisted of one soil-saprotrophic fungus and several prokaryotes including two cellulolytic bacteria. In winter, one soil-saprotrophic fungus and some prokaryotes including a cellulolytic bacterium and two bacteria involved in nitrogen cycling were identified as module hubs. One earthworm as substrate feeder (ASV68) was found to be the connector in summer, while three other connectors including one predatory myxobacterium, one saprotrophic fungus and one prokaryote were found in winter.

Connectors and module hubs are determinants of the stability of a network since they are essential in connecting the modules as well as the nodes within a module (Guimera and Nunes Amaral 2005). We found that the saprotrophic fungi and/or cellulolytic bacteria related to decomposition were among the determinants across all the seasons, suggesting decomposition as the central functional process in these high organic soils. Interestingly, eukaryotes especially some protists, played the central role in the network interactions in summer, which again suggests that eukaryotes are the important drivers of seasonal dynamics in microbiomes in fen peatlands. Protists were also found to be the central hub in the microbiome network in other soils, connecting diverse bacterial and fungal communities (Xiong et al., 2018). These findings underpin the importance of protists in driving the dynamics in belowground interactions and thus trophic interactions in soil foodwebs.

## 4. Conclusions

We investigated the seasonal dynamics in prokaryotic and eukaryotic microbiomes in three types of fen peatlands in Northern Germany. To our best knowledge, this is the first study reporting the seasonal dynamics in peat microbiomes regarding all domains of life in rewetted fen peatlands. Interestingly, the community compositions of eukaryotes (fungi, protists, and metazoa) showed significant seasonal dynamics in all sites, while the prokaryotic community composition remained stable across the seasons in almost all sites. The microbial drivers of the seasonal dynamics differed between the sites, and their majority consisted of saprotrophic and arbuscular mycorrhizal fungi, and bacterivorous protists and metazoa. The divergence of co-occurrence patterns in summer was attributed to the increased connections of protists to protists and protists to the other microbial groups. These findings suggest that microbial eukaryotes and not prokaryotes define the seasonal dynamics in microbiomes in fen peatlands, underpinning the importance of eukaryotes, especially of under-characterized protists and metazoa, in understanding the dynamics of soil foodwebs and ecosystem functionality.

## Acknowledgements

We thank Florian Beyer from the University of Rostock for providing the site maps. This study was supported by the European Social Fund (ESF) and the Ministry of Education, Science and Culture of Mecklenburg-Western Pomerania (Germany) within the scope of the project WETSCAPES (ESF/14-BM-A55-0034/16 and ESF/14-BM-A55-0030/16).

