## Supplementary for "Eukaryotic rather than prokaryotic microbiomes change over seasons in rewetted fen peatlands"

### Seasonal dynamics in prokaryotic and eukaryotic microbiomes in rewetted fen peatlands

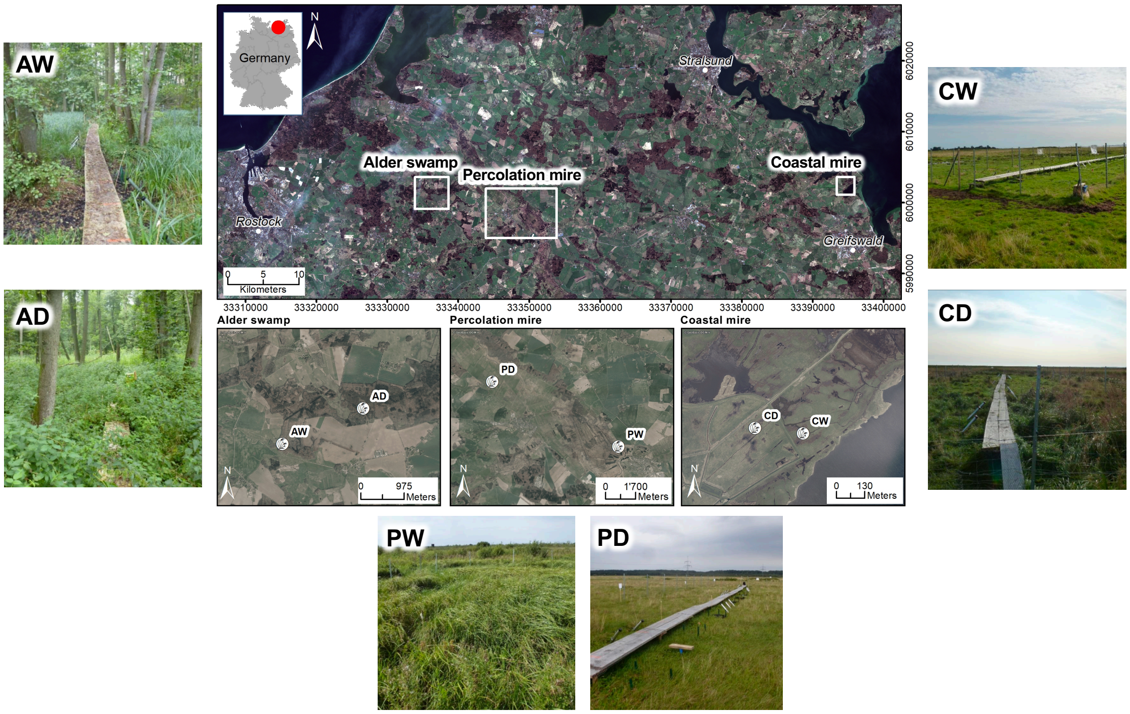


**Figure S1** Map showing locations of sampling sites: alder forest drained (AD) and rewetted (AW), coastal fen drained (CD) and rewetted (CW), and percolation fen drained (PD) and rewetted (PW). Modified from Jurasinski et al. (2020).

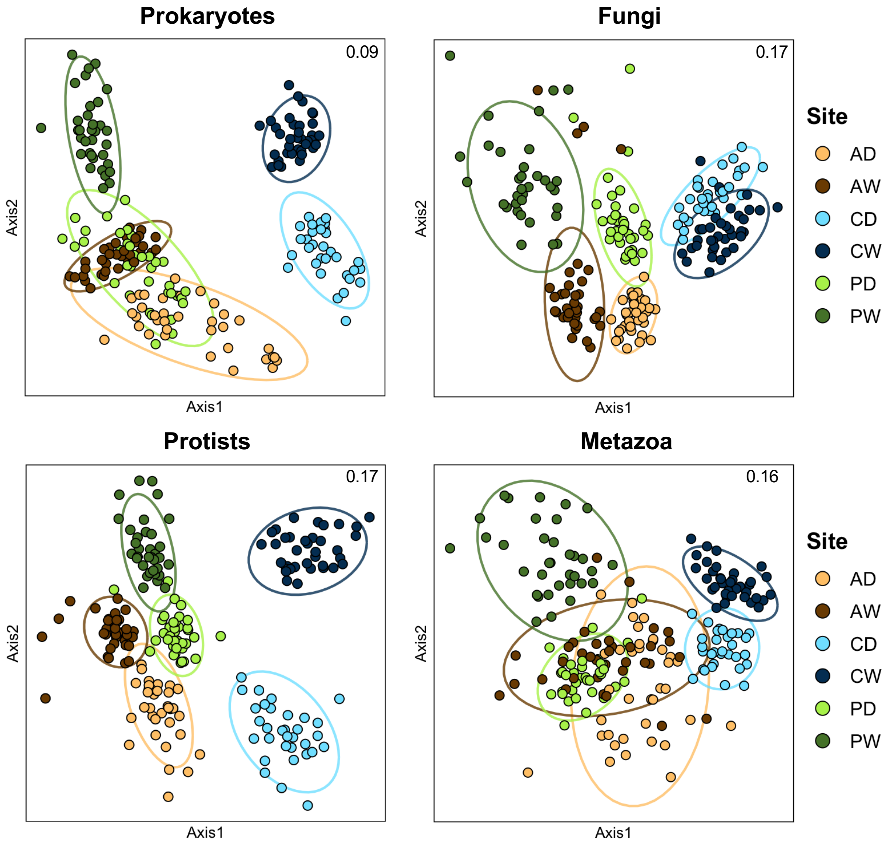


**Figure S2** NMDS plots based on the Bray-Cutis dissimilarities showing community compositions of prokaryotes and eukaryotes (fungi, protists and metazoa) in all sites. Numbers on the top right indicate the stress values.

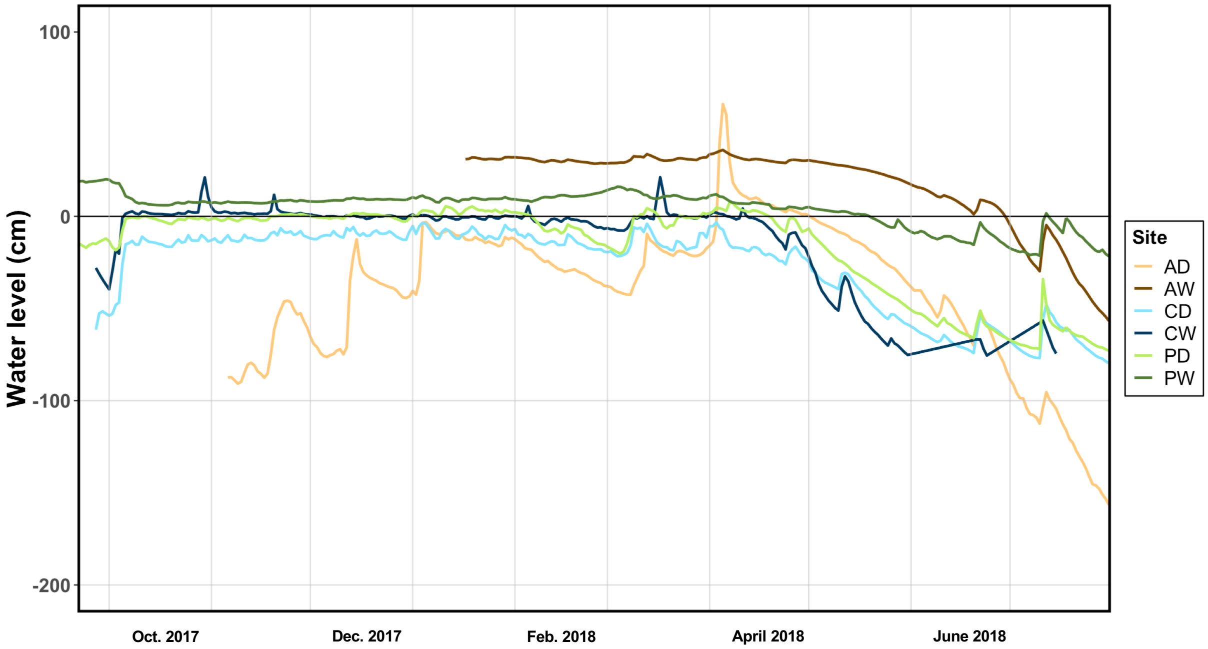


**Figure S3** Groundwater level monitored every 15 min, shown as mean value of each day from 2017-9-22 to 2018-7-31.

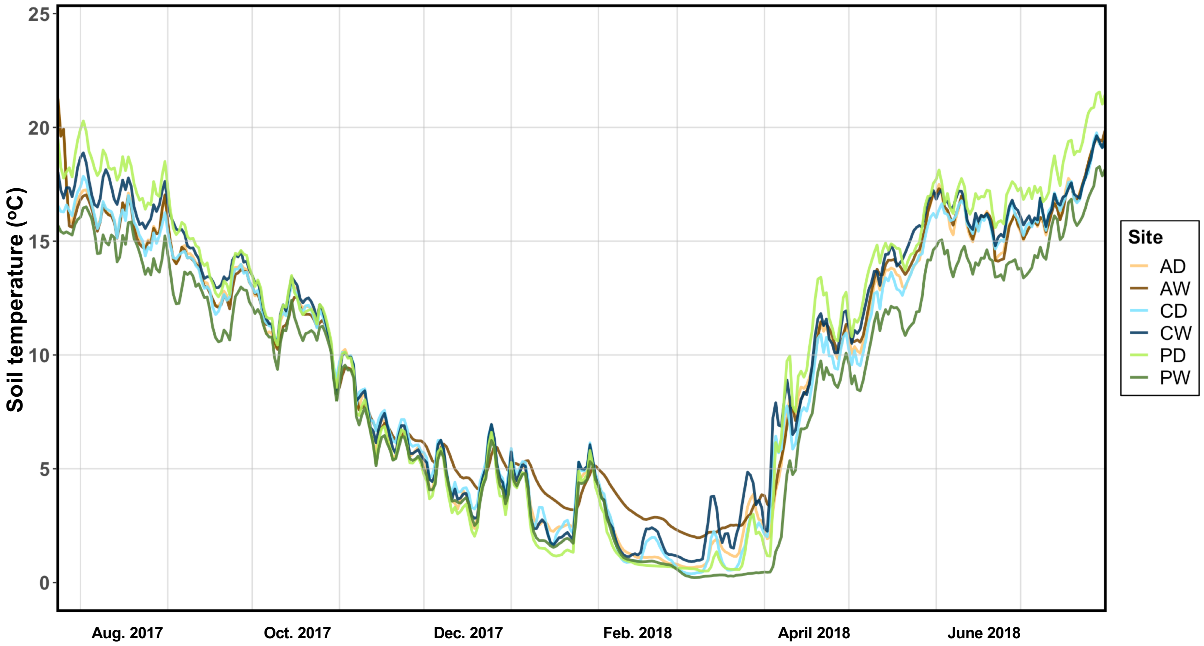


**Figure S4** Soil temperature monitored every 15 min at two depths (5 cm and 15 cm), shown as mean value of two depths and of each day from 2017-7-24 to 2018-7-31.

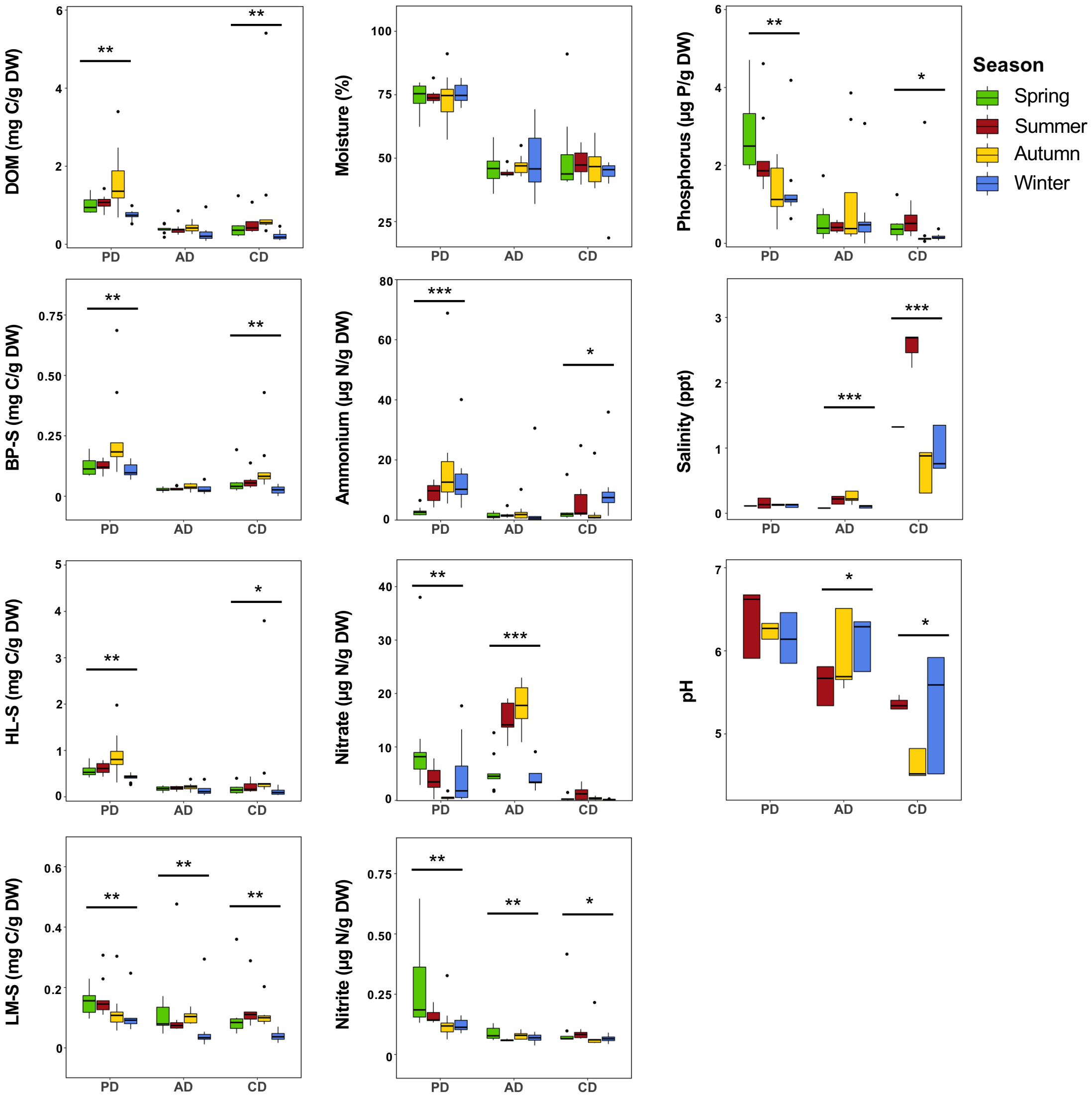


**Figure S5** Seasonal dynamics in soil edaphic properties in the three drained sites. Asterisks indicate the significant seasonal changes in one site (Kruskal-Wallis test, *adjusted $P<0.05$, **adjusted $P<0.01$, ***adjusted $P<0.001$). DOM, dissolved organic matter; BP-S, biopolymer substances; HL-S, humic-like substances; LM-S, low-molecular substances; DW, dry weight. pH in spring is not shown as a different measure method was used in spring.

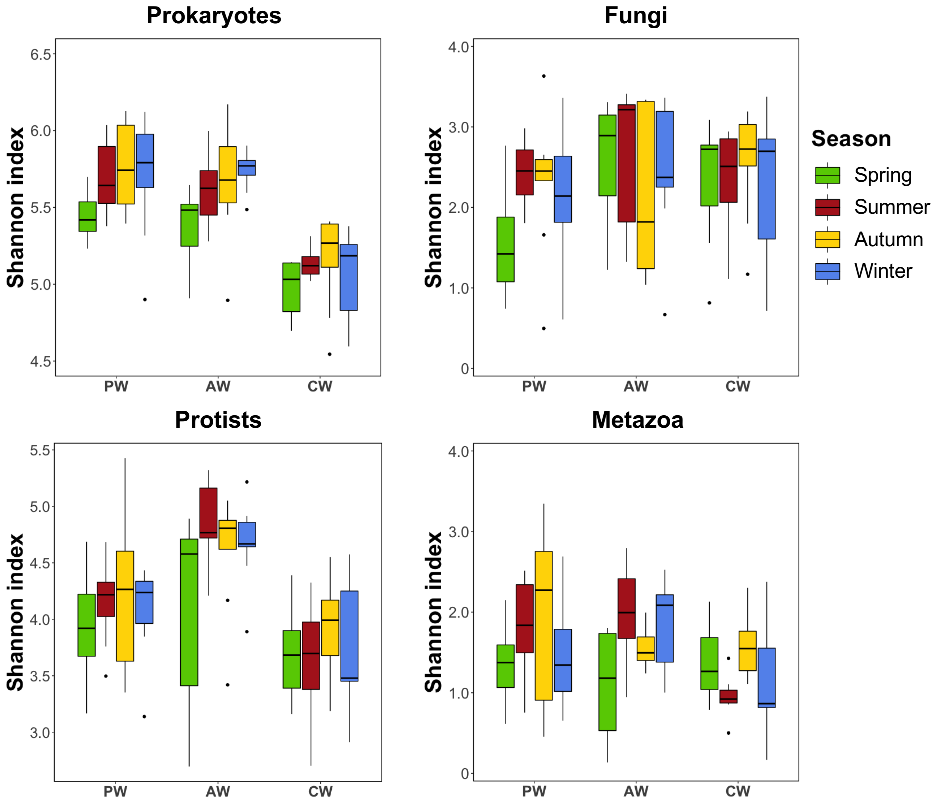


**Figure S6** Seasonal dynamics in alpha-diversity in the three rewetted sites. No significant seasonal changes were observed from any site.

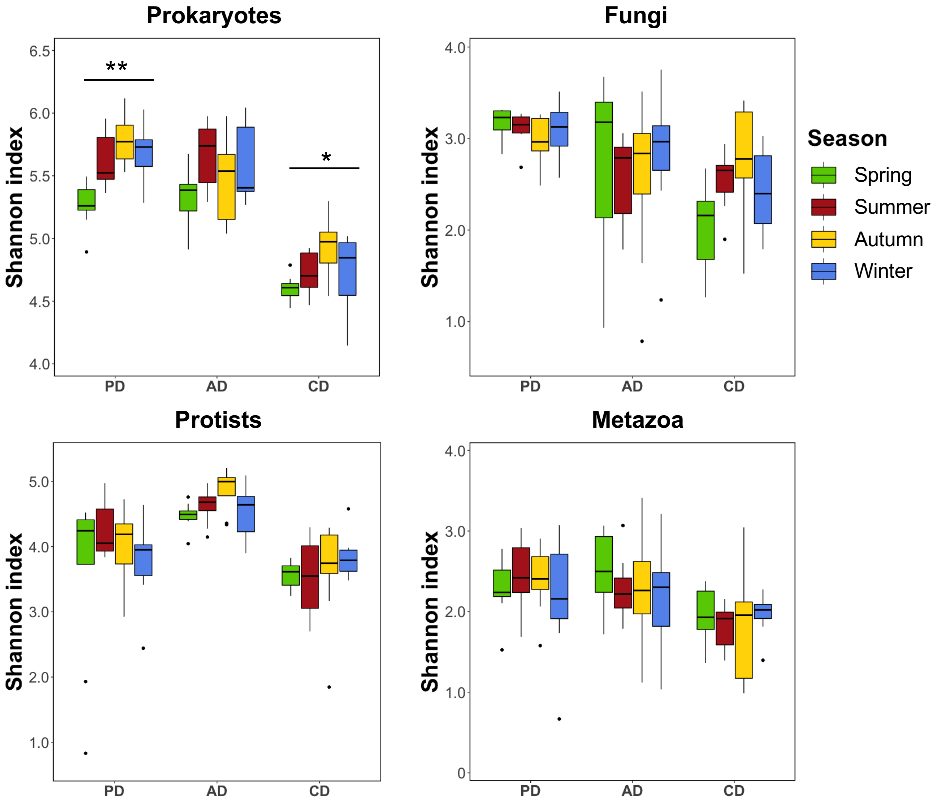


**Figure S7** Seasonal dynamics in alpha-diversity in the three drained sites. Asterisks indicate the significant seasonal changes in one site (Kruskal-Wallis test, *adjusted $P<0.05$, **adjusted $P<0.01$).

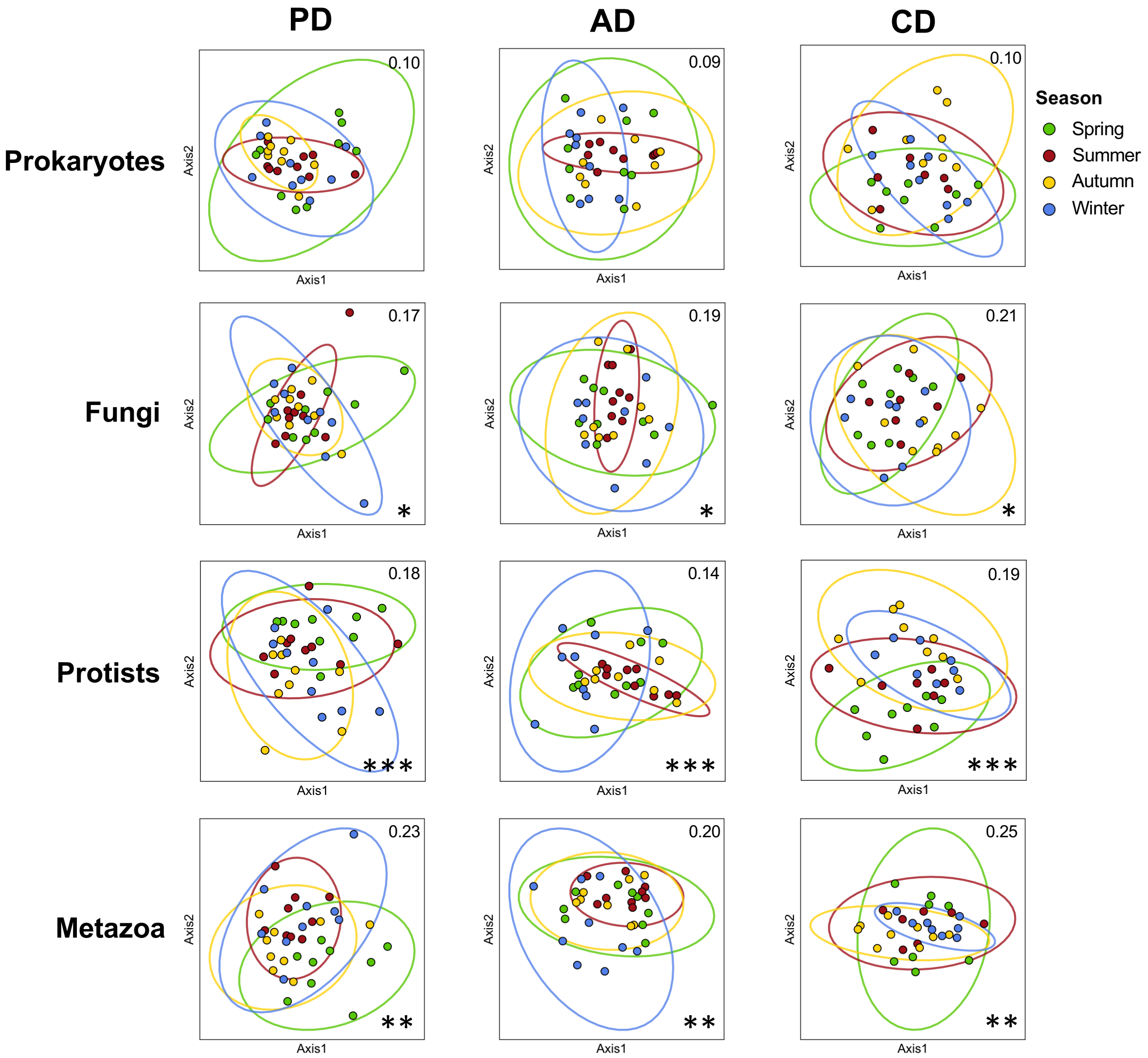


**Figure S8** NMDS plots based on the Bray-Cutis dissimilarities showing community compositions of prokaryotes and eukaryotes (fungi, protists and metazoa) in each drained site. Numbers on the top right indicate the stress values. Asterisks represent the significance level of PERMANOVA (*adjusted $P<0.05$, **adjusted $P<0.01$ or ***adjusted $P<0.001$) as indicated in Table S1.

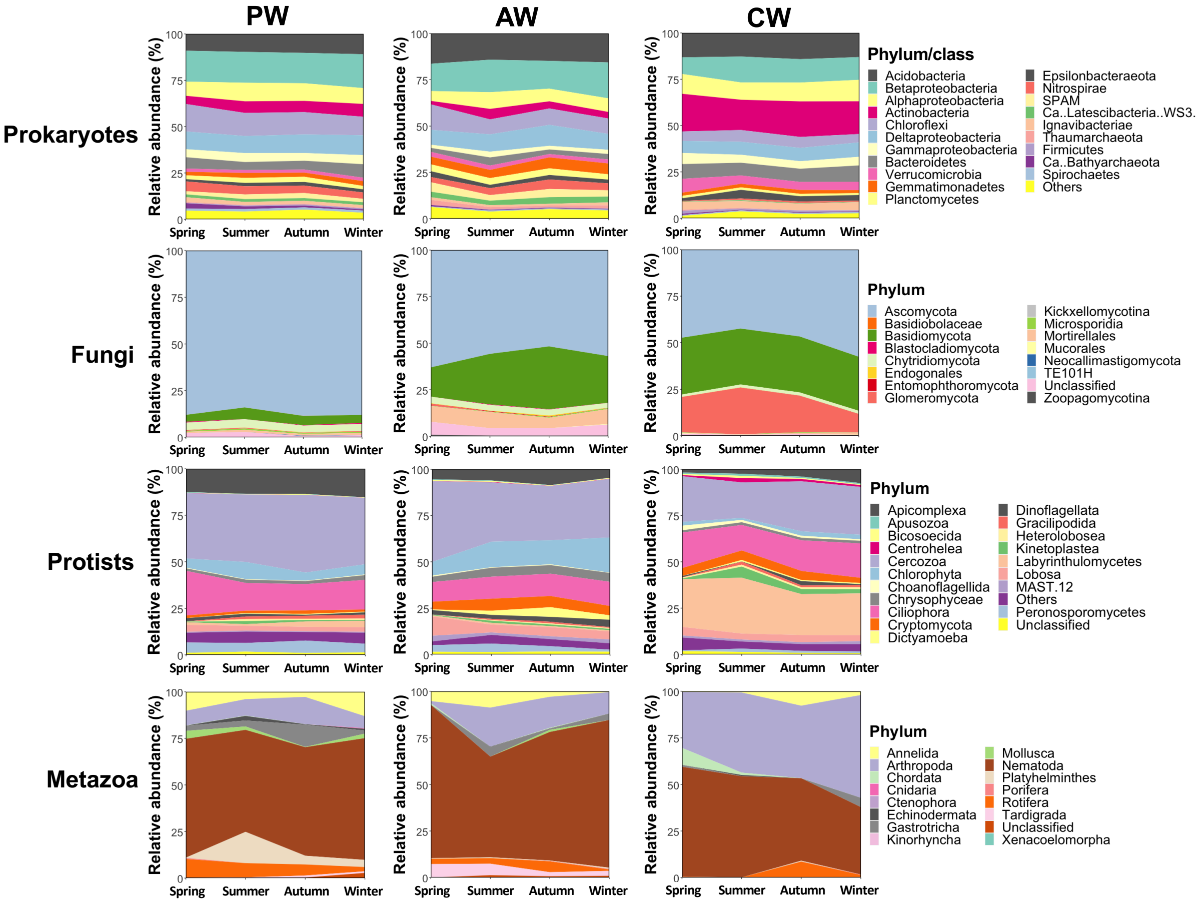


**Figure S9** Seasonal changes of pro- and eukaryotic taxa at phylum or class level in the three rewetted sites.

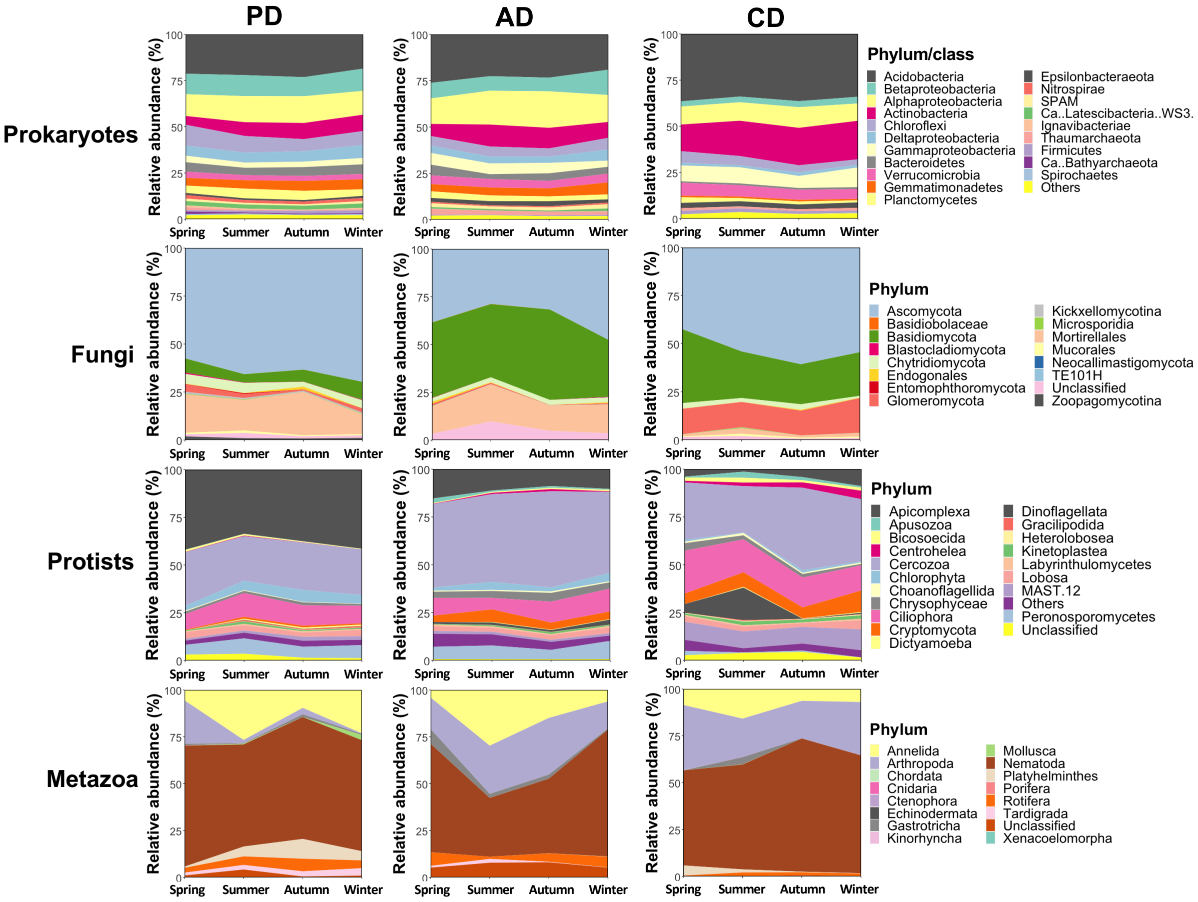


**Figure S10** Seasonal changes of pro- and eukaryotic taxa at phylum or class level in the three drained sites.

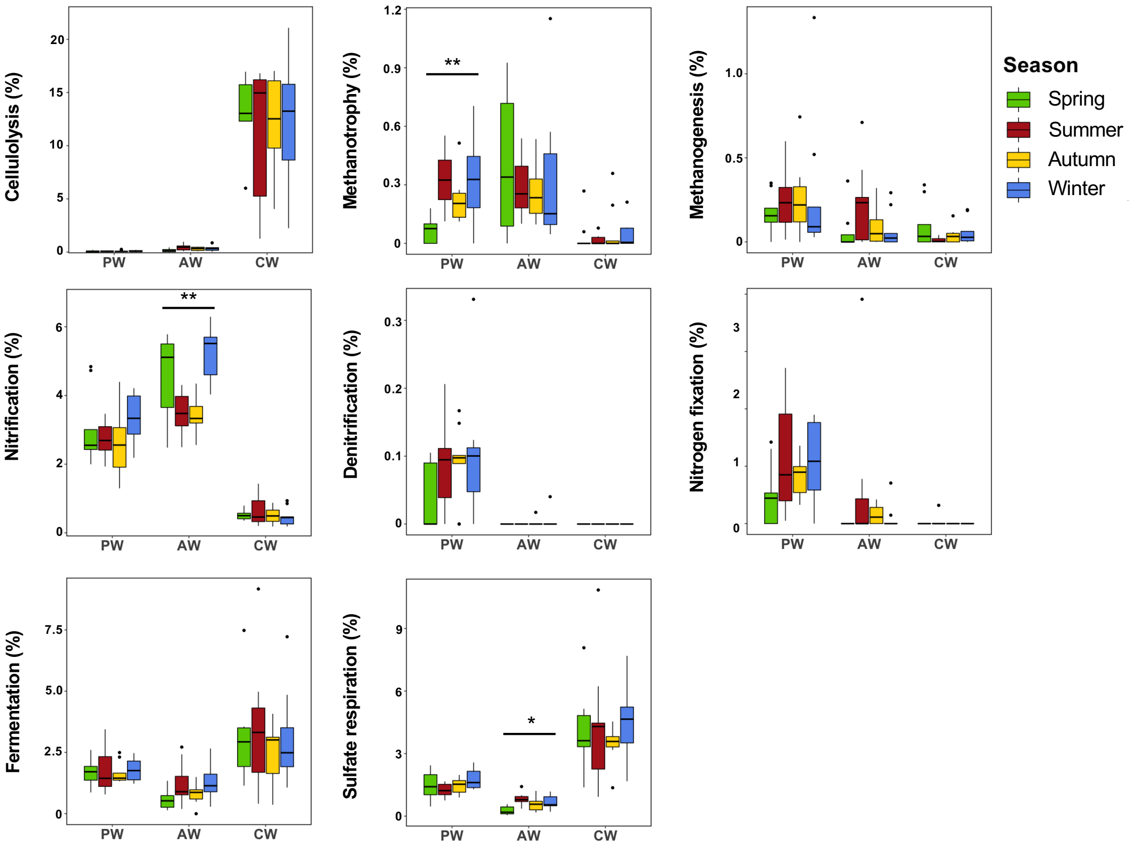


**Figure S11** Seasonal dynamics in predicted prokaryotic functions in the rewetted sites. Asterisks indicate the significant seasonal changes in one site (Kruskal-Wallis test, **adjusted *P*<0.01).

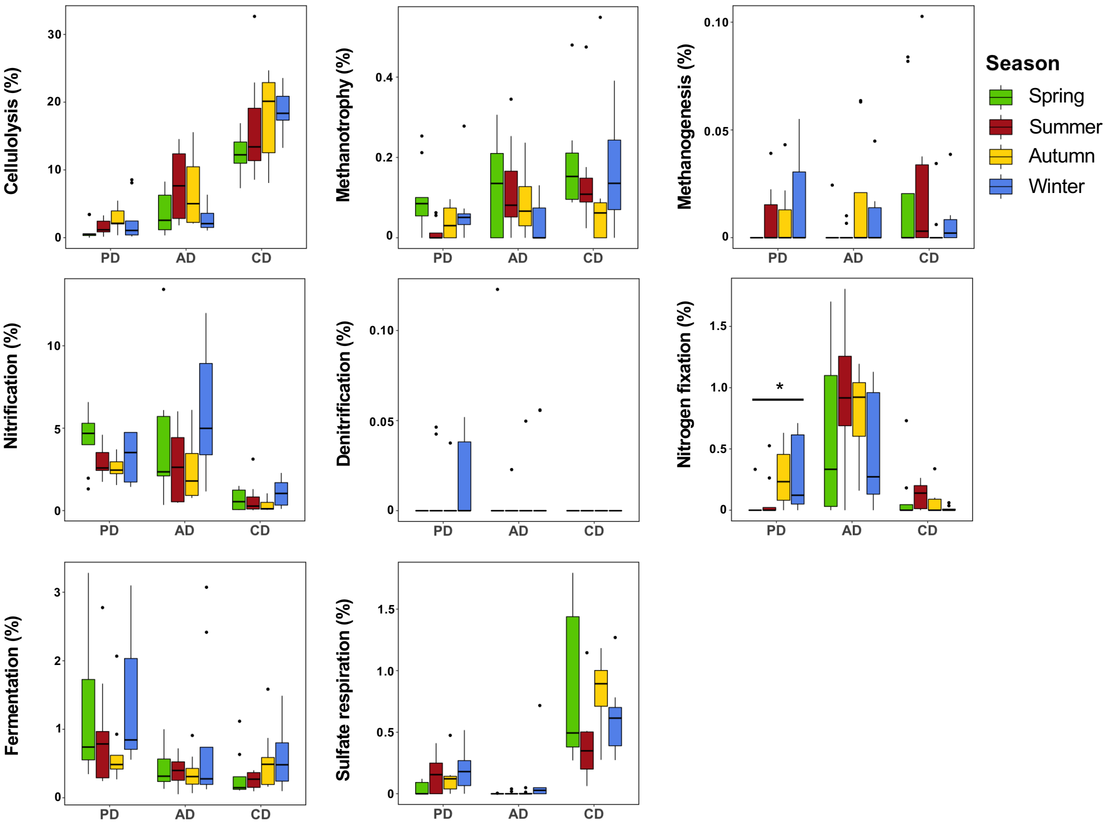


**Figure S12** Seasonal dynamics in predicted prokaryotic functions in the drained sites. Asterisks indicate the significant seasonal changes in one site (Kruskal-Wallis test, *adjusted $P$<0.05).

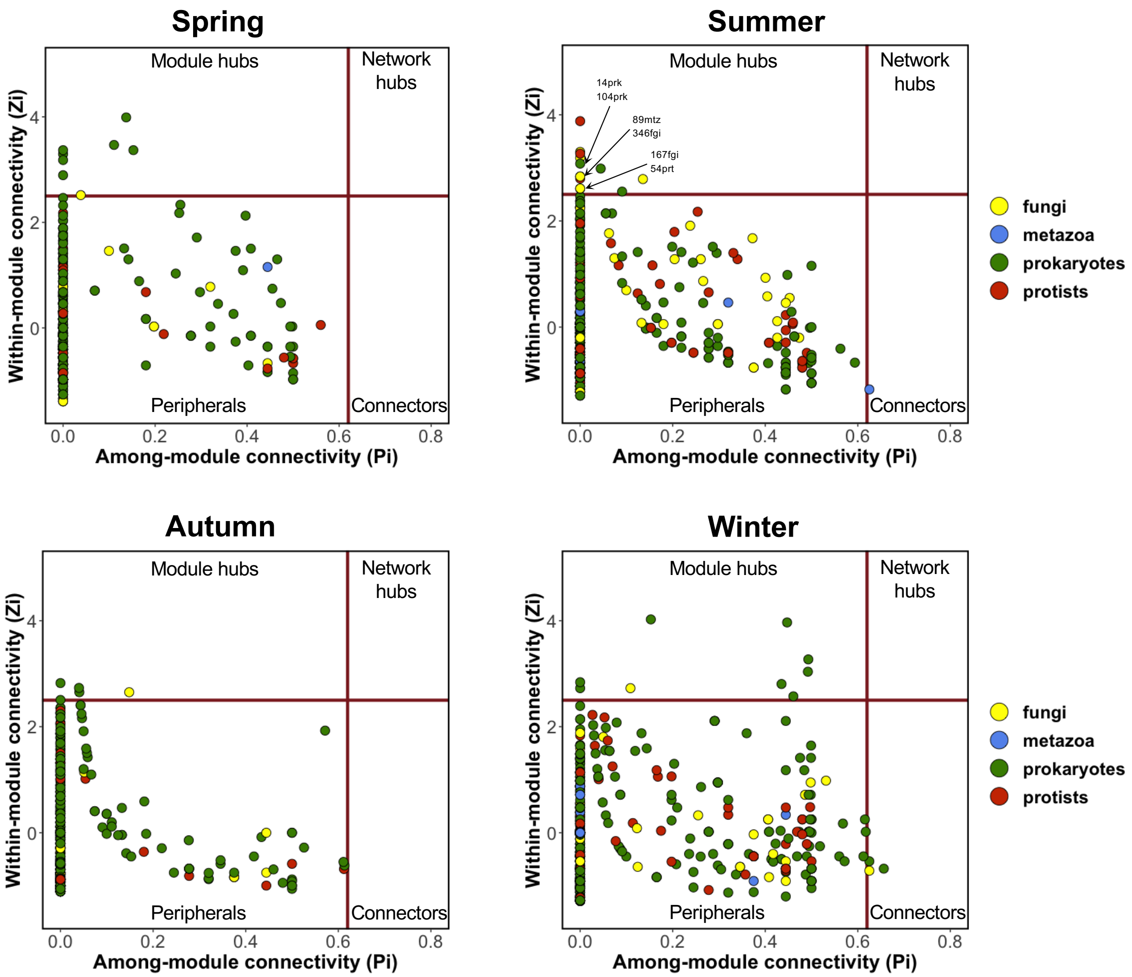


**Figure S13** Zi-Pi plots showing the topological roles of the ASVs in each season. The overlapped points are labelled separately. prk, prokaryote; fgi, fungi; prt, protist; mtz, metazoa.

**Table S1** PERMANOVA showing the significance of seasonal dynamics in community compositions in each site

| Site | Prokaryotes | | Fungi | | Protists | | Metazoa | |
|---|---|---|---|---|---|---|---|---|
| | $R^2$ | Adjusted *P* | $R^2$ | Adjusted *P* | $R^2$ | Adjusted *P* | $R^2$ | Adjusted *P* |
| PW | 0.107 | 0.176 | 0.115 | 0.035* | 0.110 | 0.013* | 0.125 | 0.012* |
| AW | 0.164 | 0.003** | 0.150 | 0.003** | 0.134 | 0.002** | 0.164 | 0.003** |
| CW | 0.124 | 0.053 | 0.128 | 0.023* | 0.134 | 0.002** | 0.161 | 0.003** |
| PD | 0.134 | 0.075 | 0.129 | 0.012* | 0.138 | 0.001*** | 0.162 | 0.003** |
| AD | 0.133 | 0.075 | 0.128 | 0.012* | 0.154 | 0.001*** | 0.139 | 0.005** |
| CD | 0.130 | 0.090 | 0.147 | 0.012* | 0.182 | 0.001*** | 0.148 | 0.005** |

Asterisks indicate the significant seasonal changes in community composition in one site (*adjusted $P < 0.05$, **adjusted $P < 0.01$ or ***adjusted $P < 0.001$ based on three rewetted or drained sites).

**Table S2** Topological features of the networks from different seasons

| | Spring | Summer | Autumn | Winter |
|---|---|---|---|---|
| Edge number | 1882 (9) | 2774 (20) | 3738 (22) | 3590 (26) |
| Vertex number | 414 | 531 | 524 | 507 |
| Average path | 5.95 | 4.75 | 5.13 | 4.72 |
| Edge density | 0.022 | 0.02 | 0.027 | 0.028 |
| Assortativity degree | 0.508 | 0.291 | 0.25 | 0.239 |
| Diameter | 17 | 13 | 12 | 16 |
| Number of modules | 14 | 14 | 13 | 19 |
| Modularity | 0.62 | 0.68 | 0.7 | 0.53 |

Edge number indicates the total number of correlations in a network, and the number in the bracket indicates the number of negative correlations. Vertex number indicates the number of vertex or ASVs in a network. Average path indicates the average path length of shortest paths between all pairs of vertices. Edge density indicates the ratio of the number of edges to the number of possible edges. Assortativity degree indicates a preference for a network's vertices to attach to others that are similar in some way. Diameter indicates the length of the longest geodesic. Modularity indicates the strength of division of a network into modules.

**Table S3** The seasonally changing ASVs and their functional traits in rewetted sites (provided in an additional file)

**Table S4** The network module hubs and connectors and their functional traits (provided in an additional file)
